# Single-cell analysis reveals the range of transcriptional states of circulating human neutrophils

**DOI:** 10.1101/2022.02.22.481522

**Authors:** Gustaf Wigerblad, Qilin Cao, Stephen Brooks, Faiza Naz, Manasi Gadkari, Kan Jiang, Sarthak Gupta, Liam O’Neil, Stefania Dell’Orso, Mariana J. Kaplan, Luis M. Franco

## Abstract

Neutrophils are the most abundant leukocytes in human blood and are essential components of innate immunity. Until recently, neutrophils were considered homogeneous and transcriptionally inactive cells, but both concepts are being challenged. To date, neutrophils have been characterized based on discrete parameters including cell-surface markers, buoyancy, maturation status, or tissue localization. Single-cell RNA sequencing (scRNA-seq) offers an unbiased view of cells along a continuum of transcriptional states. However, the use of scRNA-seq to characterize neutrophils has proven technically difficult, explaining in part the paucity of published single-cell data on neutrophils. We have found that modifications to the data analysis pipeline, rather than to the existing scRNA-seq chemistries, can significantly increase the detection of human neutrophils in scRNA-seq. We have then applied a modified pipeline to the study of human peripheral blood neutrophils. Our findings indicate that circulating human neutrophils are transcriptionally heterogeneous cells, which can be classified into one of four transcriptional clusters that are reproducible among healthy human subjects. We demonstrate that peripheral blood neutrophils shift from relatively immature (Nh0) cells, through a transitional phenotype (Nh1), into one of two endpoints defined by either relative transcriptional inactivity (Nh2) or high expression of type I interferon-inducible genes (Nh3). Transitions among states are characterized by the expression of specific transcription factors. By simultaneously measuring surface proteins and intracellular transcripts at the single-cell level, we show that these transcriptional subsets are independent of the canonical surface proteins that are commonly used to define and characterize human neutrophils. These findings provide a new view of human neutrophil heterogeneity, with potential implications for the characterization of neutrophils in health and disease.

## Introduction

The understanding of heterogeneity and plasticity in hematopoietic cells is changing rapidly. Historically, a combination of cell surface markers, transcription factors, and profiles of secreted cytokines has been employed to classify cells of similar histologic appearance and ontogeny into discrete groups. The implicit assumption of this categorization, that the resulting cell “populations” or “subsets” represent polarized and fixed states, has been questioned for the past two decades by an extensive body of evidence. This is exemplified by the recent evidence that T cells and macrophages, which have long been classified in terms of subsets, can convert from one state to another and display mixed or partial profiles ^1–3^. This evidence suggests that hematopoietic cells may be best understood along a continuum of differentiation and activation states. In this context, single-cell RNA sequencing (scRNA-seq) has been an important addition to the set of analytical tools, as cells in different states may express different sets of genes and scRNA-seq offers a less biased view of cells along a continuum of transcriptional states. Our understanding of the spectrum of cell states at baseline, in response to specific stimuli, and in disease, remains limited. The extent to which different transcriptional states correspond to older classifications based on a limited number of proteins is also unclear for most cell types.

Neutrophils are the most abundant leukocytes in human blood and essential components of the innate immune system. Until recently, they were thought to be a fairly homogeneous and transcriptionally inactive cell type, but both concepts have been convincingly challenged in recent years ^4,5^. Although human neutrophils have lower total RNA content per cell than macrophages ^6^ and other hematopoietic cell types (Supplemental Table 1), they express a broad range of genes in resting conditions ^7,8^ and their transcriptome is strongly reactive to environmental stimuli ^9–11^. Neutrophils have been characterized based on discrete parameters, including cell-surface markers, buoyancy, histologic characteristics associated with maturation status, or tissue localization. These observations have led to the emergent concept of neutrophil heterogeneity, which has been the subject of recent reviews ^4,5^. In these, it has been proposed that single-cell sequencing technology is a promising avenue for a more comprehensive and less biased characterization of neutrophil states. Recent studies in mouse models, with a limited number of human samples for comparison, have applied scRNA-seq to the study of circulating and bone marrow neutrophils, and have indeed documented the existence of a range of transcriptional states ^12,13^. Direct evidence for distinct transcriptional subsets of human neutrophils has also been provided, by our group and others, in scRNA-seq studies of sex differences in the neutrophils of healthy donors ^14^ and in patients with lung cancer^15^ or COVID-19 ^16,17^. However, given their low per-cell RNA content, scRNA-seq in neutrophils remains technically challenging, explaining in part the paucity of scRNA-seq reports describing human neutrophils compared to other hematopoietic cell types. To address this, we first evaluated the technical aspects of scRNA-seq data generation and analysis. We found that a modified pipeline is necessary for proper identification of neutrophils in scRNA-seq data. We then applied such a pipeline to the transcriptional characterization of human circulating neutrophils from multiple healthy donors at the single-cell level.

## Results

### A modified analysis pipeline is required for the adequate identification of neutrophils in single-cell RNA-seq data

The standard analysis pipeline for single-cell RNA-seq data generated with the 10X Genomics platform and Illumina short-read sequencing is implemented in the widely used Cell Ranger software ^18^. This pipeline involves grouping of the sequencing reads by their cell of origin (barcode) and RNA molecule of origin (unique molecular identifier, or UMI). This is followed by a cell-calling step, in which individual barcodes are determined to be empty (not corresponding to any cell) or to represent a captured cell. The current cell-calling algorithm employed by Cell Ranger is based on the EmptyDrops method ^19^. In the first step, the algorithm sets a threshold based on the number of UMIs associated with each barcode and those that pass this threshold are classified as cells. In the second step, a set of barcodes with low UMI counts is selected and a background model is generated. The RNA profile of each barcode that was not called as a cell in the first step is then compared against the background model and those whose profile disagrees with that of the background model are called as cells. The resulting barcodes are then output in the form of a filtered matrix of the UMI counts corresponding to each gene, in each called cell. The goal of the second step is to identify cells that may have lower RNA content than those identified in the first step.

To test the ability of this method to reliably identify human neutrophils in a mixed-cell population, we first generated single-cell RNA-seq data from a red blood cell (RBC)-depleted whole-blood sample and analyzed it with the standard Cell Ranger pipeline described above. Of the called cells in the filtered matrix, 27.2% were identified as neutrophils by an unbiased algorithm based on reference transcriptomic datasets ^20^, which was a clear underrepresentation of neutrophils in human whole blood (Figure 1A). We then visualized the frequency distribution of the number of features per barcode (genes per cell), contrasting the filtered matrix with the unfiltered matrix (Figure 1B). From this, it was clear that the filtered matrix excluded many events that were near the lower end of the distribution yet formed a peak that is distinct from the null set of events with zero or near-zero features. We hypothesized that neutrophils, having lower overall transcript abundance than other cell types, could be enriched in this excluded cell population. To test this, we modified the analysis pipeline, departing from the unfiltered matrix and lowering the threshold of genes per cell based on the observed distribution. With this modification, the proportion of cells that were identified as neutrophils rose to 58.7%, which is within the expected range of neutrophils in human whole blood (Figure 1C). Correspondingly, the proportions of other nucleated cell types (T cells, B cells, NK cells, and monocytes) either fell to, or remained within, their normal ranges in human peripheral blood, indicating a more expected representation of the cell composition of the sample. Overlaying the distribution of genes per cell of the events now identified as neutrophils on that of all events in the raw matrix indicated that a substantial proportion of the events in the distinct peak we had previously observed, in fact, correspond to neutrophils (Figure 1D). To verify the identity of the cells that were rescued by the modified analysis pipeline as neutrophils, we utilized the bioinformatic tool NeutGX ^14^, and a publicly available dataset (GEO: GSE112101) of RNA-seq data in nine primary human immune cells ^11^, to identify genes that are highly expressed in neutrophils and specific to neutrophils (*FCGR3B*) or to myeloid cells (*CSF3R, NAMPT*). The expression of these neutrophil marker genes is high in the cells that were rescued by the modified analysis pipeline and classified as neutrophils, confirming their identity (Figure 1E).

**Figure 1.**
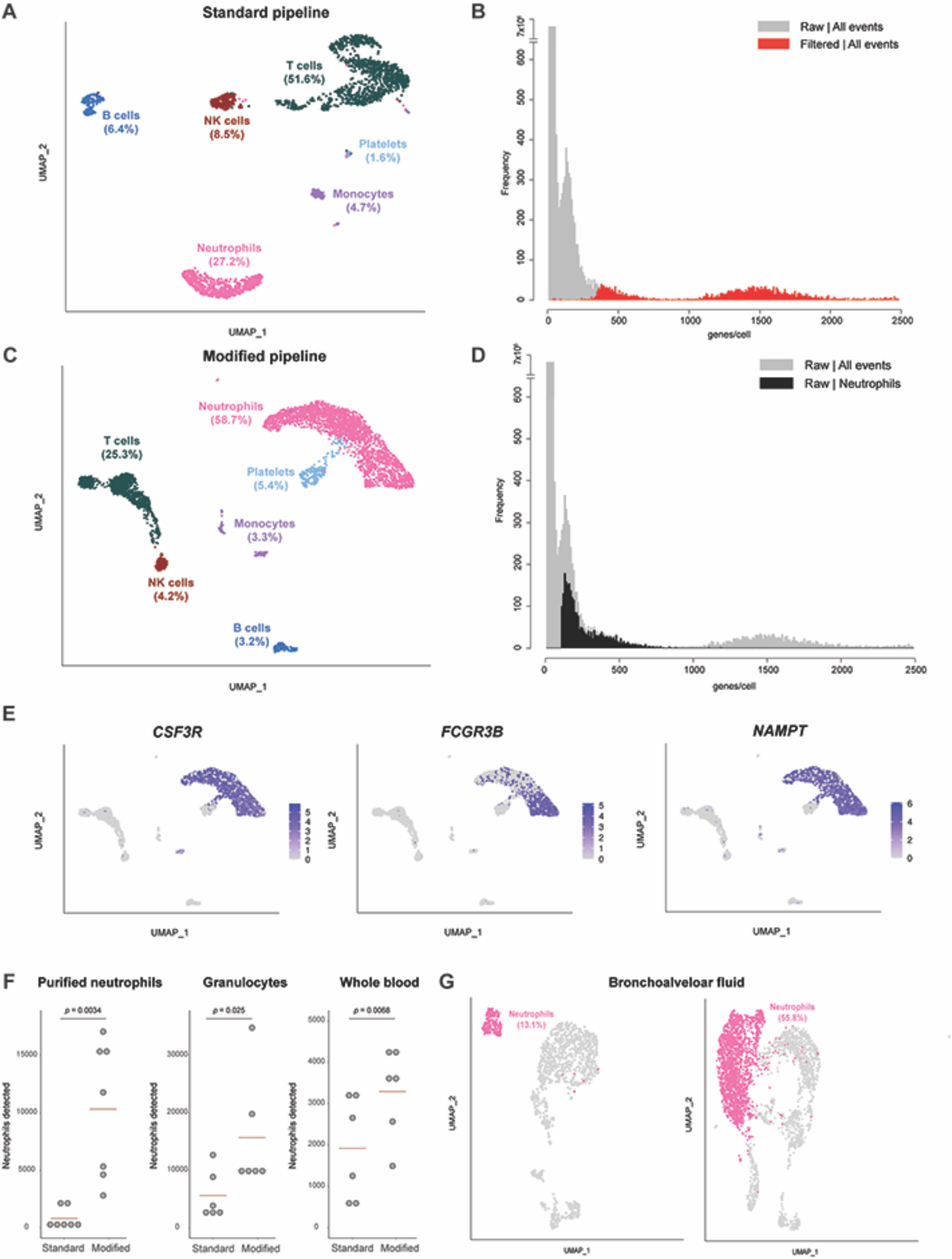
Pipeline for identification of neutrophils in scRNA-seq data. (A) Distribution of cell types identified in RBC-depleted whole-blood in a scRNA-seq analysis performed with the filtered matrix output from Cell Ranger (standard pipeline). Data from one capture are shown. (B) Frequency distribution of the number of features per barcode (genes per cell) for the dataset shown in (A), comparing data from the filtered (red) versus raw (grey) matrices. (C) Distribution of cell types identified in the dataset shown in (A) when the analysis is performed with the raw matrix output from Cell Ranger (modified pipeline). (D) Frequency distribution of the number of features per barcode (genes per cell) for the dataset shown in (A) and (C), with the distribution for cells identified as neutrophils in the analysis of the raw matrix (modified pipeline) highlighted in black. (E) Feature plot on the UMAP shown in (C) for 3 genes expected to be highly expressed in human neutrophils. (F) Number of neutrophils detected by the standard or modified pipelines in samples from the same subjects processed by three methods. Each dot represents one biological replicate (one unrelated healthy donor). Statistical testing results are from a paired t-test. (G) Proportion of neutrophils identified in a published scRNA-seq dataset of BAL fluid from patients with severe COVID-19 infection, comparing the results of the standard pipeline (left) with those of the modified pipeline (right).

In practice, depending on the requirements of specific experimental settings, human neutrophils are purified by different methods. Therefore, we systematically compared the performance of the standard or modified pipelines for neutrophil scRNA-seq in cells purified by three common methods. Whole blood from each of seven healthy donors was simultaneously processed by three methods prior to cell capture for scRNA-seq: RBC-depleted whole blood, granulocytes from density-gradient centrifugation, and immunomagnetically-purified neutrophils. In all sample types, the number of neutrophils detected was significantly higher with the modified scRNA-seq pipeline than with the standard pipeline (Figure 1F).

We then asked whether the same principle could be applied to improve the identification of neutrophils in scRNA-seq experiments with samples from other tissues. To test this, we analyzed a recently published dataset (GEO: GSE145926) of bronchoalveolar lavage (BAL) samples in patients with severe COVID-19 ^21^. With the standard analysis pipeline, 13.1% of cells were identified as neutrophils, compared to 55.8% of cells with the modified pipeline (Figure 1G).

A related approach was proposed recently online (https://support.10xgenomics.com/single-cell-gene-expression/software/pipelines/latest/tutorials/neutrophils. Accessed 9 February 2022). It involves bypassing the second step of the standard cell-calling algorithm, forcing the Cell Ranger program to call a set number of events as cells, and including intronic reads. This is followed by filtering of non-cell events based on the number of genes per cell. We performed a side-by-side comparison of this approach with our simpler, modified pipeline, and found that both are capable of rescuing neutrophils in single-cell data, and comparable in terms of the specific cells and genes identified (Supplemental Figure 1).

These results indicate that a modified analysis pipeline is required for adequate identification of neutrophils in scRNA-seq data, and that the cell-calling threshold along the frequency distribution of genes per cell is the key variable that has prevented standard analysis pipelines from identifying neutrophils.

### Human circulating neutrophils consist of distinct and reproducible transcriptional subsets

Recent observations by our group and others indicate that neutrophils from humans and mice exist in distinct transcriptional states ^12–14,16,17^. Taking advantage of the improved analysis pipeline, we directly evaluated this by performing scRNA-seq on highly pure and abundant neutrophil samples from healthy donors. To minimize the risk of potential changes in gene expression induced by gradient centrifugation, osmotic lysis of RBCs, or positive-selection antibodies, we studied neutrophils purified directly from whole blood by immunomagnetic negative selection. Flow cytometry was performed on each sample to document that the cells loaded for capture were highly pure, viable, and with no evidence of early apoptosis (Figure 2 A-C and Supplemental Table 2). As expected, the modified analysis pipeline identified a high proportion of neutrophils that would have been excluded by the standard pipeline (Figure 2D). A total of 72,183 purified circulating human neutrophils were analyzed. This analysis revealed four distinct transcriptional clusters (Fig. 2E), which were highly reproducible in samples obtained from seven unrelated healthy donors and processed independently (Figure 2F). For clarity of display and to facilitate future comparisons of our data with those from other studies in humans or other species, we have classified these clusters as Nh0 (neutrophils, human, cluster 0) through Nh3. A table with the complete set of marker genes for each cluster is provided in Supplemental Dataset 1.

**Figure 2.**
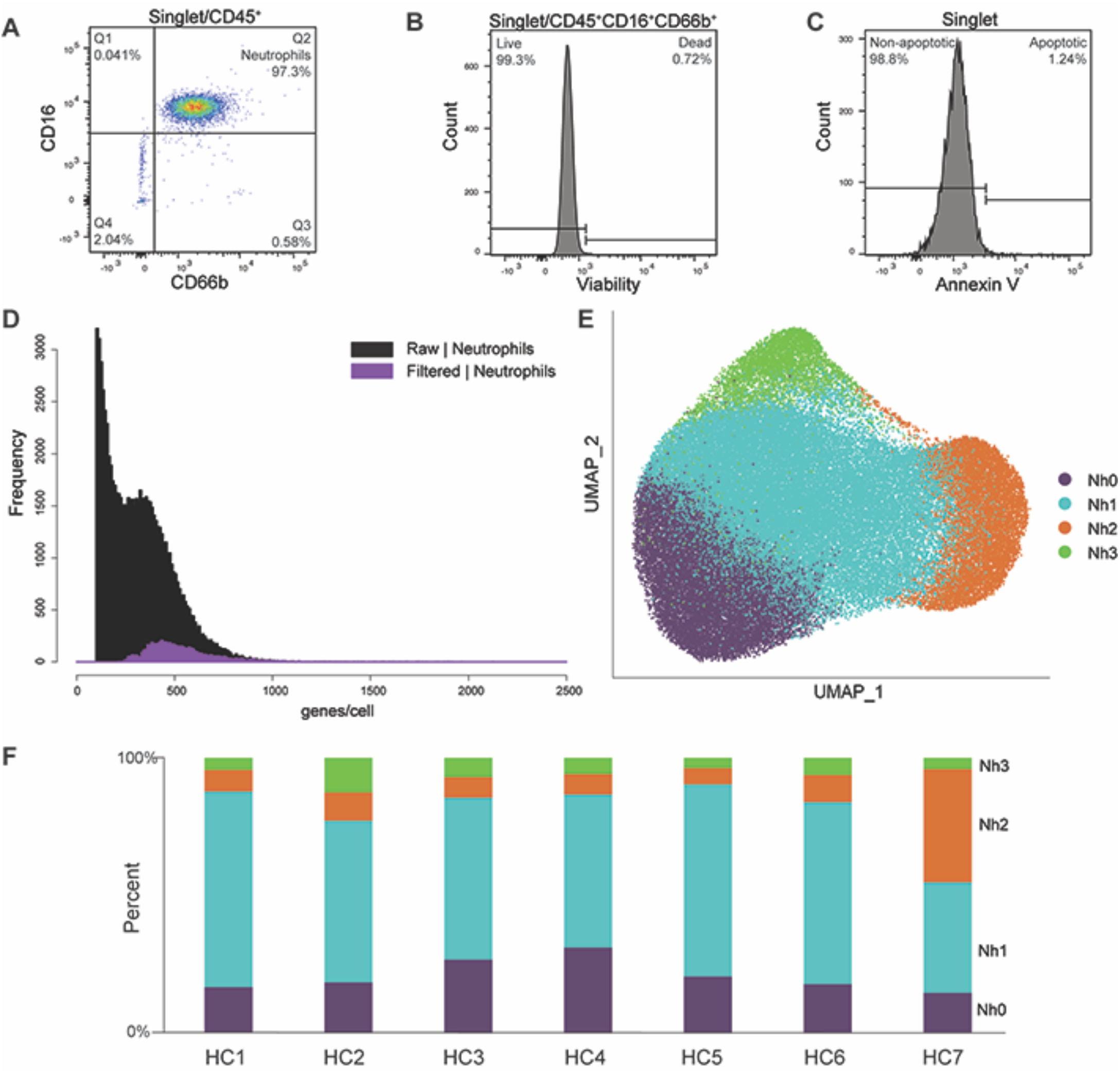
Circulating human neutrophils consist of distinct transcriptional subsets. (A-C) Flow cytometry documentation of human neutrophil purity and viability. A representative sample is shown for each panel. Purity was defined as the proportion of CD66b+CD16+ events among CD45+ events, as shown in (A). Viability was assessed by uptake of an amine-binding dye, as shown in (B). Evidence of early apoptosis was assessed by annexin-V staining, as shown in (C). Results for each sample are in Supplemental Table 2. (D) Frequency distribution of the number of features per barcode (genes/cell) in the purified neutrophils dataset, comparing data from the filtered (purple) and raw (black) matrices. (E) Two-dimensional projection (UMAP) of 72,183 purified circulating human neutrophils showing clusters Nh0 - Nh3. (F) Bar graph showing the cluster proportion of the neutrophils from each of seven healthy controls (HC1 – HC7).

Nh0 neutrophils represent approximately 20% of circulating neutrophils (mean: 22.1%, range: 14.4 – 30.1%) and are characterized by higher expression of genes that have been found to be characteristic of bone marrow neutrophils and are therefore associated with more immature neutrophil states ^8,13^. These include the genes *MMP9, ITGAM, FCN1, CAMP, CYBB, CST3*, which encode known or candidate neutrophil granule proteins ^8,22^. The genes encoding vimentin (*VIM*), thioredoxin (*TXN*), and several proteins of the S100 family (*S100A6, S100A8, S100A9, S100A11*, and *S100A12*) are also differentially expressed in Nh0 cells compared to other clusters. Of note, the gene encoding the membrane metalloendopeptidase CD10, which at the protein level is associated with more mature neutrophils, is also more highly expressed in Nh0 cells, highlighting the complementary information offered by protein- and transcript-level measurements (Figures 3A and 3B). Nh1 neutrophils represent the majority of circulating neutrophils (mean: 57.1%, range: 40.3 – 71.3%) and appear to be in a more mature state, as indicated by higher expression of the genes *AIF1, CXCR2*, and *TXNIP* (Figures 3A and 3B). Compared to other clusters, Nh1 neutrophils have a less distinct pattern of expression: contrary to other clusters, none of the top expressed genes in Nh1 are uniquely expressed in that cluster (Figure 3C). Nh2 neutrophils, which represent approximately 14% of circulating neutrophils (mean: 13.6%, range: 5.8 – 41%), are characterized by higher expression of two specific long non-coding RNAs (*MALAT1* and *NEAT1*) and of the gene encoding the G-CSF receptor (*CSF3R*), relative to other clusters (Figure 3B). Finally, Nh3 neutrophils, which correspond to approximately 7% of circulating neutrophils (mean: 7.2%, range: 3.8 – 12.6%), represent a very distinct cellular state, with substantially higher levels of expression of type I interferon (IFN)-inducible genes including *HERC5, IFI16, IFIT1, IFIT2, IFITM2, IFITM3*, and *ISG15 (*Figure 3A-D). Given that the marker genes for Nh3 neutrophils are primarily protein-coding genes expressed at very low levels in any of the other clusters, we tested whether this IFN-regulated gene-high neutrophil phenotype was also detectable at the protein level. We performed single-cell Western blotting in purified neutrophils, using antibodies recognizing ISG15 and IFITM3 (Figure 3E). We found discrete sets of neutrophils that express these proteins at high levels, and the proportion of cells in which one or both proteins is detectable is within the percentage range for Nh3 neutrophils calculated from the gene expression data (Figure 3E-F). Interestingly, most of the cells that express both of the ISG15 and IFITM3 transcripts are from the Nh3 cluster (Figure 3F), indicating the enrichment for IFN-related genes in that cluster.

**Figure 3.**
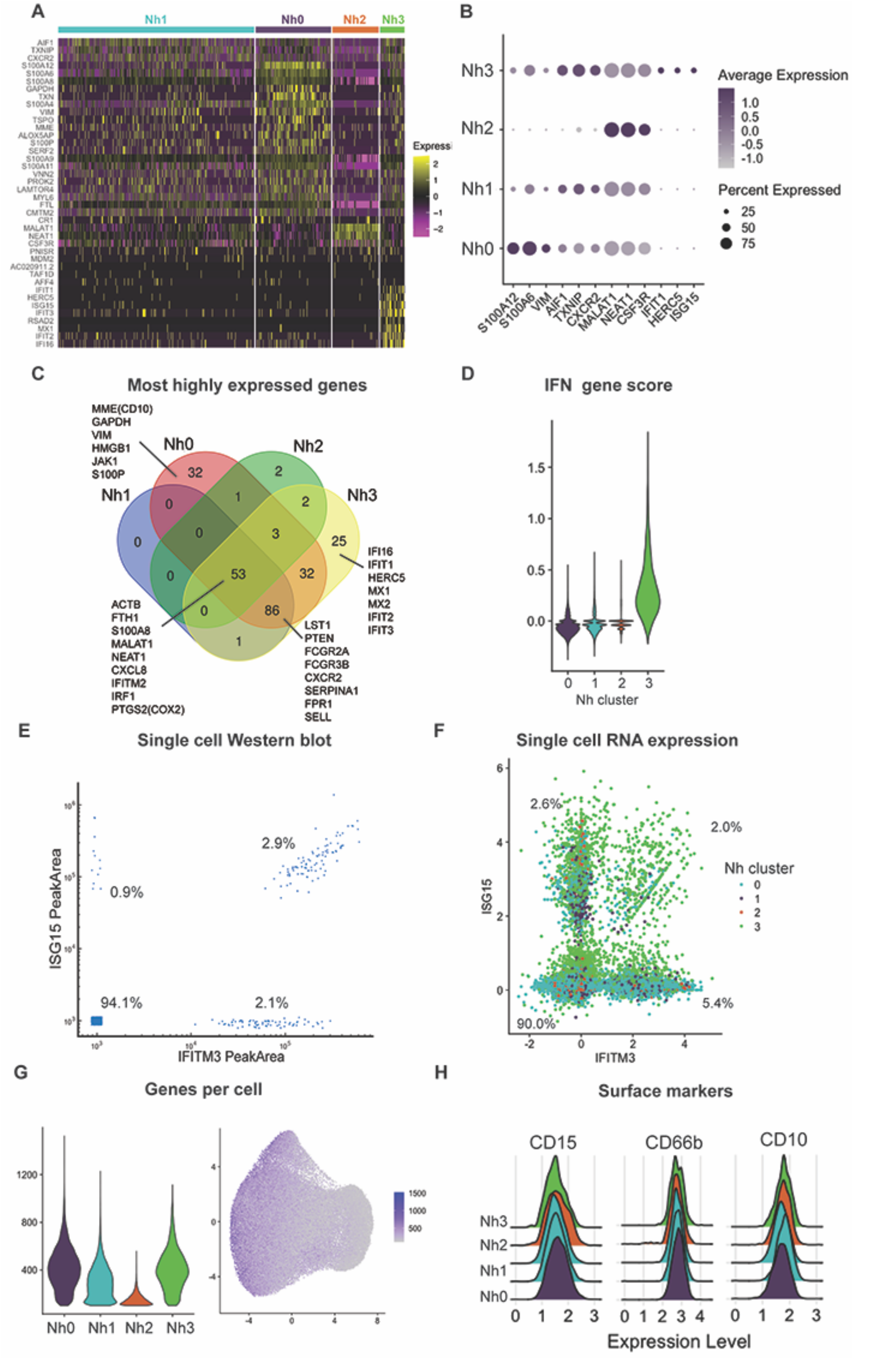
Neutrophil transcriptional subsets vary by type and number of genes expressed. (A) Heatmap of the top marker genes from each cluster. Each row represents one gene and each column represents one cell. The cells corresponding to each cluster are grouped, as indicated by the colored bars. The top marker genes were defined by their adjusted p-value and log2 (fold-difference) on differential expression analysis (expression in a cluster versus expression in all other clusters). Genes with adjusted p-value = 0 and log2FD ≥ 0.5 in any cluster are shown. (B) Dot plot of the top 3 marker genes for each neutrophil cluster, showing the average expression level and the percent of cells expressing the gene in each cluster. (C) Venn diagram displaying the intersection of the top genes in each cluster by absolute expression. (D) Violin plot showing the score per cluster for a panel of IFN-related genes, as described in ^27^. (E) Single-cell Western blot on 3,300 neutrophils, with antibodies against the proteins ISG15 and IFITM3. (F) Neutrophil single cell RNA expression of the same targets as in (F) – ISG15 and IFITM3. (G) Violin plot of the number of genes per cell in each cluster (left) and distribution of the number of genes per cell on the UMAP projection (right). (H) Ridge plots showing the distribution of CD10, CD15, and CD66b surface protein expression among cells in each transcriptional cluster. Surface expression and RNA-seq were measured simultaneously, by CITE-seq.

We compared the distribution of the number of genes detected in cells from each cluster and found that Nh2 neutrophils have a substantially lower number of genes per cell (Figure 3G). We then asked whether this difference was the result of a true biological difference between the cells in that cluster or an artifact of the clustering algorithm, whereby cells with lower read counts were classified as a distinct group. To test the latter hypothesis, we performed a down sampling analysis, in which we re-ran the entire analysis pipeline, but reducing the input reads in one of the samples by 50%. If the Nh2 cluster was in fact simply the result of cells with lower read counts being clustered together, then we would expect the down sampling of input reads to result in a higher proportion of Nh2 neutrophils. We found no change in the proportion of Nh2 neutrophils after down sampling, in the reduced sample or overall (Supplemental Figure 2), indicating that this cluster is unlikely to represent a clustering artifact driven by cells with lower read counts and instead more likely represents a distinct cluster of neutrophils with higher expression of specific genes (Figure 3B) but lower overall transcriptional output.

As with other hematopoietic cell types, neutrophils have been studied and classified almost exclusively in terms of discrete cell-surface markers measurable by flow cytometry or immunohistochemistry. To test whether the observed transcriptional subsets correlate to surface expression of one or more of the canonical proteins that have been used to characterize and group neutrophils, we performed Cellular Indexing of Transcriptomes and Epitopes by Sequencing (CITE-seq), with a custom panel of oligonucleotide-conjugated antibodies targeting CD10, CD11b, CD11c, CD14, CD15, CD16, CD24, CD33, CD35, CD45, CD66b, CD107a, CD184, and HLA-DR. This method allows simultaneous measurement of surface protein abundance and transcriptome characterization at the single-cell level ^23^. We found that the surface expression level of each of these proteins was similar among Nh0 – Nh3 neutrophils (Figure 3H and Supplemental Figure 3), indicating that these four transcriptional subsets offer a view of circulating human neutrophil heterogeneity that is independent of the canonical surface proteins that are commonly used to define and characterize these cells.

### Nh2 and Nh3 cells are endpoints in the transcriptional trajectory of human neutrophils

An important advantage of scRNA-seq is that it offers an opportunity to study cells along a range of transcriptional states, including those that fall between theoretically more stable endpoint states. This has, in turn, offered the possibility of ordering single-cell states along pseudotemporal trajectories (pseudotime), which indicate how far a given cell has moved along a continuum of biological progress. We employed the R package Monocle 3 ^24^ to construct a single-cell trajectory of circulating human neutrophils, with the immature (Nh0) neutrophils as the root. From this, it is evident that the Nh2 and Nh3 clusters represent distinct endpoints in the transcriptional trajectory of circulating neutrophils, while the Nh1 cluster represents an intermediate state (Figure 4A). We then looked for genes that vary between clusters of circulating neutrophils and grouped these into modules that have a similar pattern of expression. This identified five modules of co-expressed genes (Figure 4B), which we mapped back to the trajectory map. Module 1 genes are most highly expressed in the immature (Nh0) neutrophil cluster, module 3 genes in the NEAT1/MALAT1 (Nh2) neutrophil cluster, and module 5 genes in the IFN (Nh3) neutrophil cluster. The genes in modules 2 and 4 are more highly expressed in the transitional (Nh1) neutrophil cluster, but they represent distinct regions along the trajectory: module 2 genes appear to characterize a transitional state between Nh0 and Nh2 neutrophils, whereas module 4 genes characterize a transitional state between Nh0 and Nh3 neutrophils (Figure 4C).

**Figure 4.**
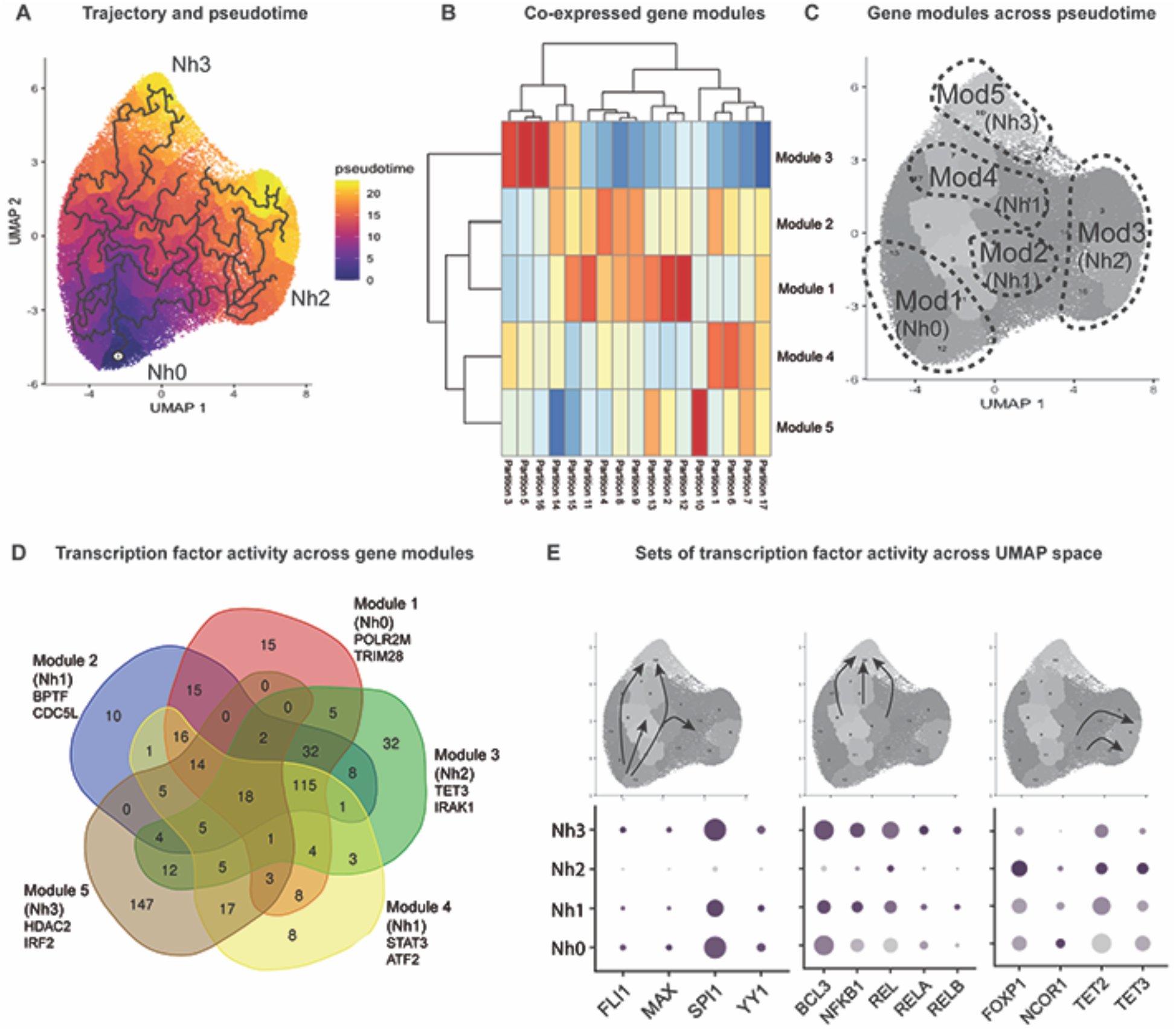
Nh2 and Nh3 cells are endpoints in the transcriptional trajectory of circulating human neutrophils. (A) Trajectory analysis showing the learned graph on the UMAP space with the pseudotime ordering by color. (B) Heatmap showing unsupervised classification of genes that vary across clusters of circulating neutrophils into five clusters of co-expressed genes. (C) Correspondence between the five modules of co-expressed genes and the four transcriptional clusters of circulating human neutrophils. (D) Venn diagram of transcription factors associated with cis-regulatory elements most likely to regulate the co-expressed genes in each module. Gene lists from the modules were used as input in binding analysis for regulation of transcription. The overlap across modules for the top-ranking transcription factors (Irwin-Hall p-value < 0.01) is shown. (E) Transcription factor gene expression changes along the transcriptional trajectory of circulating human neutrophils. Three patterns are shown: transcription factors expressed along the Nh0-Nh1-Nh3 trajectory, but not in Nh2 cells (left); transcription factors expressed in the transition from Nh1 to Nh3 cells (middle), and transcription factors expressed in the transition from Nh1 to Nh2 cells(right).

Modules of co-expressed genes offer an opportunity to infer common transcriptional regulatory elements, without the assumptions and potential biases inherent to inference based on known functions or on genomic localization with respect to other genes or to DNA sequence motifs. To infer candidate transcription factors that regulate the sets of co-expressed genes in each neutrophil module, we employed binding analysis for regulation of transcription (BART), a method that relies on experimental evidence of protein-DNA interactions for over 400 known transcription factors across a variety of cell types ^25^. We then selected the transcription factors associated with cis-regulatory elements most likely to regulate the co-expressed genes from each module (Irwin-Hall p-value < 0.01) and compared these across modules. The modules corresponding to Nh2 and Nh3 neutrophils have the highest number of predicted transcription factors uniquely associated with them (Fig. 4D). The transcription factors at the intersections of neutrophil modules are also informative, as the expression of genes encoding specific transcription factors varies along the transition from one neutrophil cluster to another. For example, the genes encoding the transcription factors FLI1, MAX, SPI1, and YY1 are expressed along the trajectory from the immature (Nh0) to the IFN (Nh3) states, but not in the MALAT1/NEAT1 (Nh2) neutrophil cluster (Fig. 4E, left). Similarly, the transition from the intermediate (Nh1) cluster to the IFN (Nh3) cluster, is marked by increased expression of genes involved in NFkB signaling (Fig. 4E, center). In contrast, the transition from the intermediate (Nh1) cluster to the NEAT1/MALAT1 (Nh2) cluster is characterized by increased expression of the genes encoding the transcriptional repressor FOXP1 and the methylcytosine dioxygenases TET2 and TET3.

Our results support a model in which Nh2 and Nh3 cells represent endpoints in the transcriptional trajectory of circulating human neutrophils. Distinct sets of transcription factors, at least some of which are regulated at the level of transcription, orchestrate the transition from a less mature state (Nh0 cells) to one endpoint state or the other, via an intermediate state (Nh1 cells) that corresponds to the majority of circulating neutrophils.

## Discussion

Our findings indicate that circulating human neutrophils are transcriptionally heterogeneous cells, which can be classified based on their transcriptional state into one of four clusters (Nh0-Nh3) that are highly reproducible among healthy human subjects. We demonstrate that neutrophils transition transcriptionally from relatively immature (Nh0) cells, through an intermediate phenotype (Nh1), into one of two endpoints defined by either relative transcriptional inactivity (Nh2) or higher expression of IFN-induced genes (Nh3). More broadly, our findings demonstrate the feasibility of applying scRNA-seq to the study of human neutrophils obtained by different methods, by means of a modified analysis pipeline that significantly improves the identification of neutrophils in scRNA-seq datasets.

Recent studies have applied scRNA-seq to the study of murine neutrophil development in states of health or experimental infection ^12,13^, and have found clear evidence of neutrophil transcriptional heterogeneity. One of these studies also analyzed CD33+ cells sorted from whole blood from a human donor ^12^, while the other analyzed a publicly available scRNA-seq dataset generated from human bone marrow neutrophils as part of the Human Cell Atlas ^13^, suggesting that human neutrophils also exhibit distinct patterns of transcriptional heterogeneity. Our group and others have also provided recent evidence for transcriptional subsets of human neutrophils in scRNA-seq studies of sex differences in neutrophils obtained from healthy donors ^14^ and in patients with lung cancer ^15^ or COVID-19 ^16,17^. However, due to their lower RNA content relative to other cell types, scRNA-seq with human neutrophils remains technically challenging and not well standardized, and it is common in human scRNA-seq studies for neutrophils to be missing or drastically under-represented with respect to their expected proportions ^17,21,26,27^. One possibility is that nucleases or proteases in neutrophil granules could interfere with the standard cell capture, cell lysis, or library preparation steps in scRNA-seq. However, after testing several modifications to the standard 10X chemistry, we did not find a clear benefit to the addition of nuclease or protease inhibitors. Another possibility is that the standard cell-calling algorithms that are routinely used by most labs are not optimal for the differentiation of neutrophils from the background distribution of empty capture beads, thus excluding most neutrophils from downstream analyses. We found this to be the most likely source of neutrophil underrepresentation and describe an alternative approach to data analysis that departs from the raw matrix of UMIs associated with each barcode and considers the observed frequency distribution of features per barcode (genes per cell). This simple modification to the analysis pipeline significantly increases the inclusion of cells that, based on their transcriptional profile, clearly represent neutrophils.

We applied the modified analysis pipeline to the study of human neutrophils enriched by immunomagnetic negative selection to very high levels of purity and viability and without evidence of early apoptosis. We analyzed 72,183 cells and found that circulating human neutrophils can be consistently clustered into four distinct transcriptional states, which we have classified as Nh0 – Nh3. The global pattern of gene expression in Nh0 cells is similar to what has been described in bone marrow neutrophils,^8,13^ with higher relative expression of various granule proteins and of several members of the S100 family. Trajectory analysis indicates that circulating neutrophils develop from this relatively immature state into a transitional cluster, Nh1, which is transcriptionally the least distinct cluster and accounts for a majority of the captured cells (∼ 60%). From this cluster, the developmental trajectory diverges towards one of two endpoint states: the Nh2 and Nh3 phenotypes. Nh2 cells are characterized by higher relative expression of specific non-coding (*NEAT1, MALAT1*) or coding (*CSF3R*) RNAs, but have a lower overall transcriptional output than other neutrophils. Accordingly, they also have higher expression of genes encoding active regulatory elements that are associated with epigenetic modulation of transcription in neutrophil development, including *TET2* and *NELFA* ^28^. Additionally, the gene encoding the transcription factor SPI1 (PU.1) which is a central factor in myeloid development ^29^, is highly expressed in all clusters except Nh2. This endpoint, therefore, likely represents the mature and transcriptionally quiescent state that has been classically associated with all circulating neutrophils. The IFN-gene-expressing Nh3 cluster is transcriptionally quite distinct from the Nh2 state. Nh3 cells express more genes, they have increased expression of IFN-inducible genes that are not significantly expressed by any other neutrophil cluster and, based on our results, their expression of key regulatory transcription factors is also distinct. The transition from Nh1 to Nh3 is associated with increased expression of genes in the NFkB family of transcription factors, which are known to play a role in the regulation of neutrophil activation, apoptosis, and NADPH oxidase activity ^30–32^. The existence of a subset of circulating neutrophils that expresses increased levels of IFN-inducible genes is now a well-validated finding in mouse and human ^12–17^, and we had previously shown that there are gender differences in the expression of the genes in this cluster ^14^. Our single-cell Western blot results indicate that this cluster is also likely to be detectable at the protein level. It is still unknown whether these cells represent neutrophils that have encountered a specific stimulus *in vivo* or if they are epigenetically committed from a precursor state. In either case, the fact that the proportion of this cluster is relatively stable among healthy donors suggests that they represent a steady state rather than an incidental finding related to a recent exposure. More studies in humans will be necessary, but data from *E. coli*-challenged mice suggest that the equivalent cluster of IFN-high neutrophils might have different bone marrow precursors than other neutrophils ^12^.

The limitations of this work can be considered in two categories. First, there are limitations related to the current state of scRNA-seq technology and data analysis methods. As with all available scRNA-seq technologies, we rely on a very shallow sampling of the transcriptome of any given cell (50,000 reads per cell in our case, but in many studies half of that or less). Data analysis methods in scRNA-seq also rely on linear (principal components analysis) and non-linear (UMAP or t-SNE) reductions from a high-dimensional ambient space into two-dimensional representations, with inevitable loss of potentially important relations between cells. The choice of clustering algorithms and parameters can also drastically affect the results, which highlights the need for standardized methods and clear reporting. Second, there are limitations related to the scope of our experiments. We focus on a single scRNA-seq chemistry (10X Genomics) which, although highly prevalent, is not the only one available. The extent to which our modified analysis pipeline can be extrapolated or adapted to other chemistries remains to be determined. Our study is also limited to circulating human neutrophils, which are of obvious biological importance but represent a minority of total neutrophils. Finally, the transcriptional subsets we describe appear to be offering a view of neutrophil heterogeneity that is independent of the very limited one afforded so far by a small set of cell-surface markers. There is, at this time, no reliable way to sort neutrophils based on transcriptional signatures while preserving viability. Therefore, experimental characterization of possible functional differences among the transcriptional subsets we have described is an important future goal.

Based on our results, we propose that human circulating neutrophils are transcriptionally dynamic cells that develop from a less mature state into one of two distinct transcriptional phenotypes that cannot be defined by common surface markers. We also propose that a modified analysis pipeline is necessary for proper representation of neutrophils in scRNA-seq studies. We hope that these findings will pave the way for better representation of neutrophils in scRNA-seq studies, to a better understanding of neutrophil heterogeneity, and to additional studies exploring the behavior of these transcriptional neutrophil subsets over time (circadian variation or variation over the human lifespan), in response to environmental or pharmacological stimuli, or under different pathologic conditions.

## Methods

### Cell purification

Human venous peripheral blood samples from healthy donors were obtained from the Department of Transfusion Medicine at the National Institutes of Health Clinical Center. For neutrophil purification, whole blood samples were collected in vacutainer glass blood collection tubes with acid citrate dextrose (ACD). Neutrophils were isolated with the EasySep Direct Human Neutrophil Isolation Kit (STEMCELL Technologies; cat. no. 19666).

For granulocyte purification, whole blood was collected in heparinized tubes. Granulocytes were isolated by dextran sedimentation of RBC pellets as previously described ^33^. Briefly, cells were first layered on a Ficoll/Hypaque gradient (GE Healthcare; cat. no. 17144003). The granulocyte/RBC fraction was then enriched by dextran sedimentation followed by RBC lysis using hypotonic solution. Granulocytes were then washed with phosphate-buffered saline (PBS). For white blood cell purification, whole blood samples were collected in heparinized tubes. White blood cells were isolated with the Erythroclear Red Blood Cell Depletion Reagent Kit (STEMCELL Technologies; cat. no. 01738).

### Documentation of cell purity and viability

Flow cytometry was used to assess the purity and viability of purified neutrophils. The cells were stained with a panel of monoclonal antibodies containing: ECD CD16 clone 3G8 (Beckman Coulter; cat.no. A33098), BV711 CD45 clone HI30 (BD Biosciences; cat.no. 564357) and FITC CD66b clone G10F5 (Biolegend; cat.no. 305104). The LIVE/DEAD Fixable Dead Cell Stain Kit with aqua fluorescent reactive dye (ThermoFisher Scientific; cat.no. L34957) was used to assess cell viability. PE Annexin V (BioLegend; cat.no. 640908) was used to assess early apoptosis activity. UltraComp eBeads Compensation Beads (ThermoFisher Scientific; cat.no. 01-2222-42) were used to perform spectral compensation. Data was collected by a BD Biosiences FACSCelesta flow cytometer, and later analyzed with Flowjo software (v10). The purity of neutrophils was specifically defined by cell-lineage markers, as the proportion of CD66b^+^CD16^+^ events among CD45^+^ events.

### Single-cell RNA-seq

From each cell purification sample, approximately 50,000 cells were centrifuged at 300g for 5 minutes at 4°C and washed twice with PBS with 0.02% bovine serum albumin (BSA). To obtain single-cell gel beads-in-emulsion (GEMs), we resuspended cells at a concentration of 1000 cells/µL and added 1µl of RNase Inhibitor (Invitrogen, Cat. N.10777-019) before loading the mix on a Chromium Comptroller Instrument (10x Genomics). Single-cell cDNAs and libraries were prepared with a Chromium Single Cell 3′ Library & Gel Bead Kit v3.1 (10x Genomics; cat. no. 1000121). Briefly, GEM-RT incubation was performed in a C1000 Touch Thermal cycler with 96-Deep Well Reaction Module (Bio-Rad; cat. no. 1851197): 53°C for 45 min, 85°C for 5 min, held at 4°C. Single-strand cDNAs were purified with DynaBeads MyOne Silane Beads (Thermo Fisher Scientific; cat. no. 37002D) and amplified with the C1000 Touch Thermal cycler with 96-Deep Well Reaction Module: 98°C for 3 min; 13 cycles of 98°C for 15 sec, 63°C for 20 s, and 72°C for 1 min; 72°C for 1 min; held at 4°C. Amplified cDNA products were cleaned with 0.6X DynaBeads MyOne Silane Beads (Thermo Fisher Scientific; cat. no. 37002D). Quality and quantity of the cDNAs were assessed on a 4200 Tape Station (Agilent Technologies) with High Sensitivity D5000 DNA Screen Tape (Agilent; cat. no. 5067-5592). The final material was amplified as follows: 98°C for 45 sec; 16 cycles of 98°C for 20 sec, 54°C for 30 sec, 72°C for 20 sec; 72°C for 1 min; held at 4°C. Libraries were diluted to the same molarity and pooled for sequencing on a NextSeq500 (Illumina) or NovaSeq6000 (Illumina) sequencers. Sequencing read lengths were 28bp for read 1, 8bp for the i7 index, and 91 bp for read 2.

Protease and RNase activity are known to be highly active in neutrophils ^34^. It should be noted, however, that the addition of a protease inhibitor (Thermo Fisher Scientific, cat. no. A32963) or an RNase Inhibitor (Ambion, cat. no. AM2682), individually or in combination, to the standard 10X protocol for cell capture and library preparation, did not increase the final cDNA concentration at the end of the library construction phase of the protocol.

### CITE-seq

TotalSeq-B oligonucleotide-conjugated antibodies (Biolegend), compatible with the 10X Genomics 3’ scRNA-seq chemistry, were used according to the manufacturer’s protocol. The panel for common markers of circulating neutrophils included antibodies targeting CD45, CD14, CD33, CD11c, CD10, CD16, CD107a, HLA-DR, CD11b, CD66b, CD35, CD24, CD184, and CD15.

### Processing and analysis of single-cell RNA-seq data

Illumina run folders were demultiplexed and converted to FASTQ format with Cell Ranger mkfastq version 4.0.0 and Illumina bcl2fastq version 2.20. Reads were further counted and analyzed with Cell Ranger count version 4.0.0 and the refdata-gex-GRCh38-2020-A reference, to generate raw and filtered matrix files.

Matrix files were imported into the R package Seurat version 4.0.1 ^35^ for downstream processing. From the raw matrices, cells with a gene number between 100-2500 and a mitochondrial gene proportion < 0.1 were selected for downstream analysis. The matrices were then normalized by the LogNormalize method. The FindVariableFeatures() function was used to select the top 2,000 variable genes, with the vst selection method. Scaling was performed by the function ScaleData() regressing out the mitochondrial gene content. Principal component analysis (PCA) and clustering were then performed on the scaled data. UMAP (version 0.2.7.0) was utilized for visualization and SingleR (version 1.4.1) was used for cell identification.

After neutrophils were identified in the dataset corresponding to each sample, they were integrated. First, contaminants were removed if they had gene expression values >1 for three marker genes specific to RBCs (HBA2), T cells (CD3G), and cells with a predominance of ribosomal RNA (RPS8). Second, genes that were shared among all datasets were identified for downstream integration. Anchors were identified with the FindIntegrationAnchors() function, and these anchors were used to integrate the neutrophils together with the function IntegrateData(). Finally, UMAP was performed on the top 8 principal components from the integrated data, and the resolution was set to 0.2 for visualization of the four clusters identified.

To examine the effect of sequencing depth on clustering and other downstream analyses, selected single samples were down sampled by 50%. Typical single-cell Illumina runs consisted of two lanes of a flow cell sequencing the same pooled libraries. 50% down sampling was accomplished by analyzing the data from a single lane.

To study neutrophil cell-state trajectories, we used the analysis toolkit Monocle3, which is implemented as an R package (version 0.2.3.0) ^24^. A principal graph was learned on the UMAP projection of the cells with the learn_graph() function. To generate a pseudotime axis, the cells were then ordered with the order_cells() function. To identify genes that vary between groups of cells in UMAP space, we used the graph_test() function; this function employs the spatial autocorrelation analysis statistic Moran’s I, which has been shown to be effective for identifying genes that vary in scRNA-seq datasets ^36^. The genes found to be variable were then grouped into modules with the function find_gene_modules(), which employs the Louvain method for community detection ^37^, to identify clusters of genes with a similar pattern of expression. To infer which transcriptional regulators are active in the cells, the module gene lists were used as input for the Binding Analysis for Regulation of Transcription (BART) pipeline ^25^. Transcription factors associated with cis-regulatory elements most likely to regulate the input gene lists (Irwin-Hall p-value <0.01) were used for further analysis with the Ghent University Bioinformatics and Evolutionary Genomics custom Venn diagram tool (http://bioinformatics.psb.ugent.be/webtools/Venn/).

### Single Cell Western Bloting

Purified neutrophils were loaded on the scWest chip (ProteinSimple), allowed to settle for 20 min, and treated according to manufacturer’s instructions. Briefly, the chip was placed in the Milo instrument (ProteinSimple) for 15 s lysis, 45 s separation, and 4 min UV exposure. The chip was then probed using antibodies against the proteins ISG15 (Cell Signaling, cat 2758) and GAPDH (Cell Signaling, cat 5174), labeled with Alexa 488/Alexa 594, and scanned in an array scanner (Molecular Devices). Chips were then stripped and reprobed with antibodies against IFITM3 (Cell Signaling, cat 59212) and rescanned. Analysis of the images was done in Scout software (ProteinSimple), where GAPDH was used as loading control cell marker and data presented as % of total cells positive for ISG15 and/or IFITM3. A total of 3300 neutrophils were used from 2 different donors.

## Supporting information

suppl data marker genes

suppl data raw expression

## Acknowledgements

This work was supported by the Intramural Research Program of the National Institute of Arthritis and Musculoskeletal and Skin Diseases (NIAMS) at the National Institutes of Health (NIH). L.M.F. also receives research support from the Division of Intramural Research at the National Institute of Allergy and Infectious Diseases (NIAID) at the NIH. This study used the high-performance computing clusters of the NIAID Office of Cyber Infrastructure and Computational Biology and of the NIAMS Office of Science and Technology. We thank Thomas Lewis at the NIH Clinical Center’s Department of Transfusion Medicine for support in the obtention of human samples. We thank James Simone at the Flow Cytometry Section of the NIAMS Office of Science and Technology, for his technical expertise and assistance.

## Author Contributions/Contributor Role Taxonomy (CRediT) Statement

Gustaf Wigerblad: Conceptualization, Data Curation, Formal analysis, Investigation, Visualization, Writing – Original Draft. Qilin Cao: Conceptualization, Data Curation, Formal analysis, Investigation, Visualization, Writing – Original Draft. Stephen Brooks: Data Curation, Formal analysis, Methodology. Faiza Naz: Investigation. Manasi Gadkari: Investigation. Kan Jiang: Data Curation, Formal analysis. Liam O’Neil: Conceptualization, Investigation. Stefania Dell’Orso: Conceptualization, Investigation. Mariana J. Kaplan: Conceptualization, Supervision, Writing – Review & Editing. Luis M. Franco: Conceptualization, Formal analysis, Supervision, Writing – Original Draft.

