## Supplementary material for "Single-cell analysis reveals the range of transcriptional states of circulating human neutrophils": Suppl figures and tables

### Supplemental Materials for
### Single-cell RNA-seq reveals distinct subsets of circulating human neutrophils

Gustaf Wigerblad, Qilin Cao, Stephen Brooks, Faiza Naz, Manasi Gadkari, Kan Jiang, Sarthak Gupta, Liam O'Neil, Stefania Dell'Orso, Mariana J. Kaplan, Luis M. Franco

**Contents:**

Supplemental Table 1. RNA content in neutrophils versus other hematopoietic cell types.

Supplemental Table 2. Purity, viability, and early apoptosis of purified neutrophils by subject.

Supplemental Figure 1. Comparison of two modified analysis pipelines for identification of neutrophils in scRNA-seq data.

Supplemental Figure 2. Down sampling analysis.

Supplemental Figure 3. Cellular Indexing of Transcriptomes and Epitopes by Sequencing (CITE-seq), with a custom panel of oligonucleotide-conjugated antibodies targeting 14 canonical neutrophil surface proteins.

Supplemental Dataset 1: Marker genes by neutrophil cluster.

Supplemental Dataset 2: Expressed genes by neutrophil cluster.

| Cell type | RNA (µg per million cells) |
|---|---|
| **Neutrophils** | 0.33320272 |
| **B lymphocytes** | 1.16230159 |
| **Monocytes** | 2.5489899 |
| **NK cells** | 0.87908497 |
| **CD4+ T cells** | 1.26430396 |
| **CD8+ T cells** | 1.39197031 |

**Supplemental Table 1**. **RNA content in neutrophils versus other human hematopoietic cell types.** Cells were purified by immunomagnetic enrichment for the specific cell subset with EasySep Human cell enrichment kits (STEMCELL Technologies). B lymphocytes, NK, CD4+ and CD8+ T lymphocytes were isolated from PBMCs by negative selection (STEMCELL Technologies; cat. nos. 19054, 19055, 19052 and 19053, respectively). Monocytes were isolated from PBMCs by positive selection (STEMCELL Technologies; cat. no. 17858). Neutrophils were isolated from whole blood by negative selection immunomagnetic purification with the EasySep Direct Human Neutrophil Isolation Kit (STEMCELL Technologies; cat. no. 19666). RNA quantity was measured on a Qubit 2.0 fluorometer (Thermo Fisher Scientific; cat. no. Q32866), with RNA BR quantitation assays (Thermo Fisher Scientific; cat. no. Q10211). RNA yield in µg per million cells represents the mean of four biological replicates (unrelated healthy donors) for neutrophils and three biological replicates for all the other cell types.

| Subject | Purity | Viability | Early Apoptosis |
|---|---|---|---|
| **HC1** | 97.3% | 99.3% | 1.2% |
| **HC2** | 98.5% | 99.5% | 2.7% |
| **HC3** | 98.0% | 99.3% | 2.0% |
| **HC4** | 95.8% | 97.4% | 1.9% |
| **HC5** | 98.2% | 99.2% | 0.4% |
| **HC6** | 97.6% | 98.2% | 1.7% |
| **HC7** | 99.6% | 99.8% | 2.6% |
| **Mean** | 97.9% | 99.0% | 1.8% |
| **Standard deviation** | 1.2% | 0.8% | 0.8% |

**Supplemental Table 2. Purity, viability, and early apoptosis of purified neutrophils by subject.** Each purified neutrophil sample was stained with antibodies against CD16, CD45, and CD66b antibodies for purity; Live/Dead fixable dye for viability, and Annexin V for early apoptosis. Flow cytometry was performed on a BD Biosiences FACSCelesta flow cytometer. The purity of neutrophils was defined as the proportion of $CD66b^{+}CD16^{+}$events among $CD45^{+}$ singlet events.

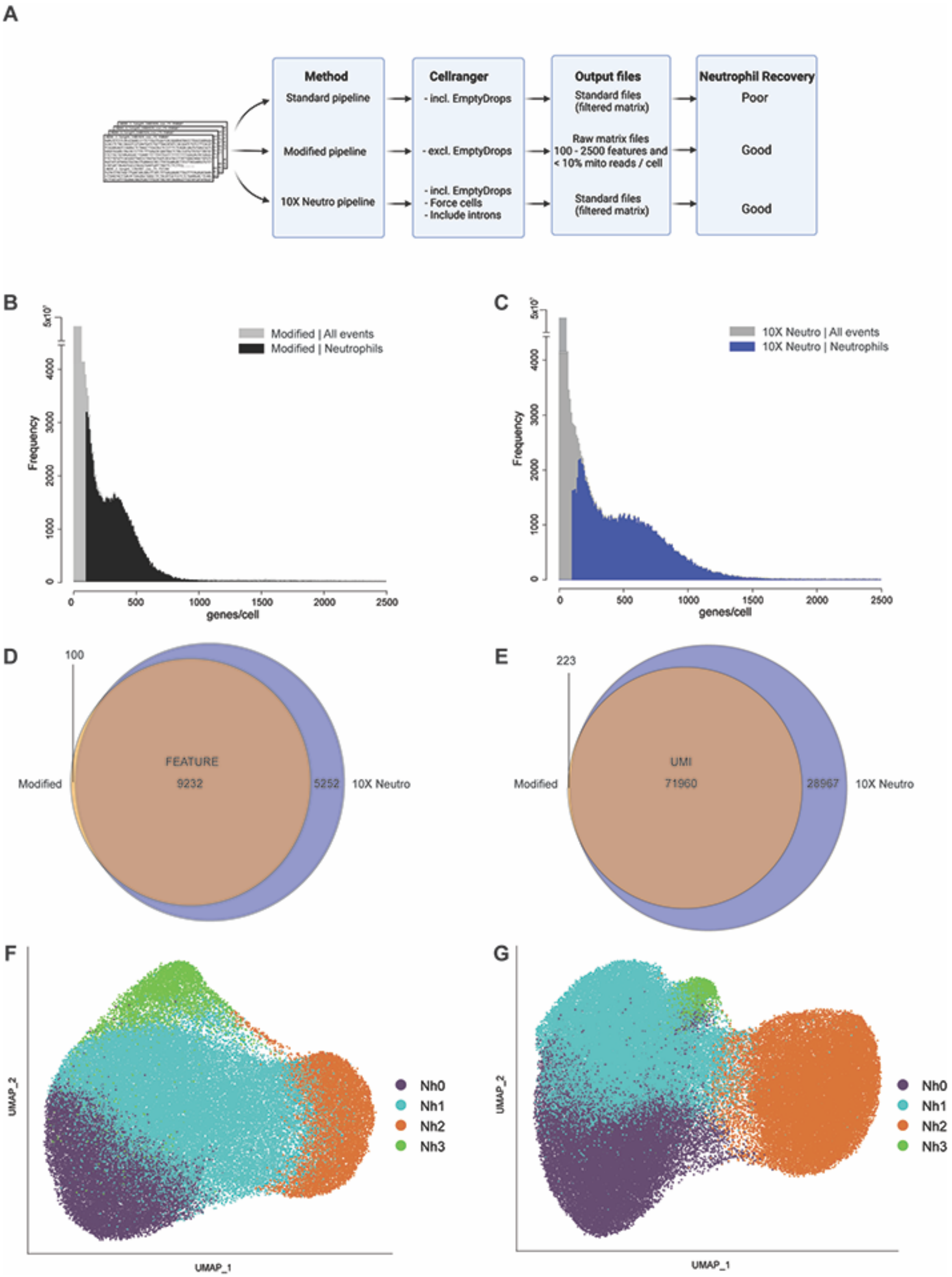


**Supplemental Figure 1**. **Comparison of two modified analysis pipelines for identification of neutrophils in scRNA-seq data.** (A) Overview of the three analysis pipelines: standard, modified, and 10X Neutro. (B) Frequency distribution of the number of features per barcode (genes/cell) in the purified neutrophils dataset processed by modified pipeline 1 (grey), with the distribution for cells identified as neutrophils (black). (C) Frequency distribution of the number of features per barcode (genes/cell) in the purified neutrophils dataset processed by modified pipeline 2 (dark grey), with the distribution for cells identified as neutrophils (blue). (D-E) Comparisons of the features and UMIs between modified pipeline 1 and modified pipeline 2. (F-G) Two-dimensional projection (UMAP) of purified circulating human neutrophils identified by modified pipeline and 10X Neutro pipeline, respectively.

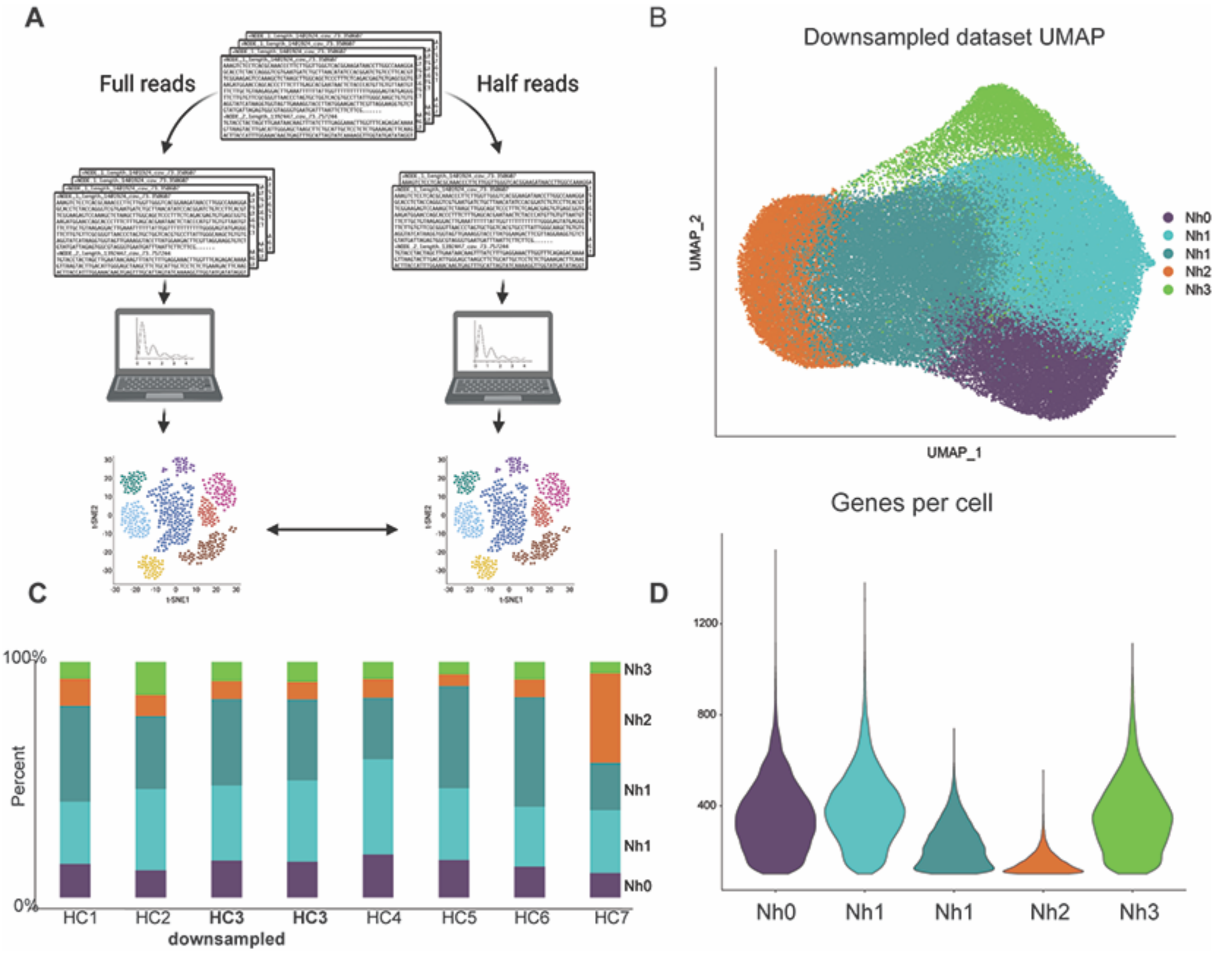


**Supplemental Figure 2**. **Down sampling analysis**. (A) Overview of the analysis pipeline. FASTQ files from one of the samples were randomly reduced by 50% and run through the entire analysis pipeline. Results were then compared with the full dataset to test whether the Nh2 cluster was simply a reflection of cells with lower read counts. (B) UMAP projection of the reduced-reads dataset, with cluster names corresponding to those in the full-read dataset (Nh0-Nh3). (C) Bar graph showing cluster proportions for all samples, including the down sampled HC3. (D) Violin plot showing the number of genes per cell in each cluster of the reduced-reads dataset.

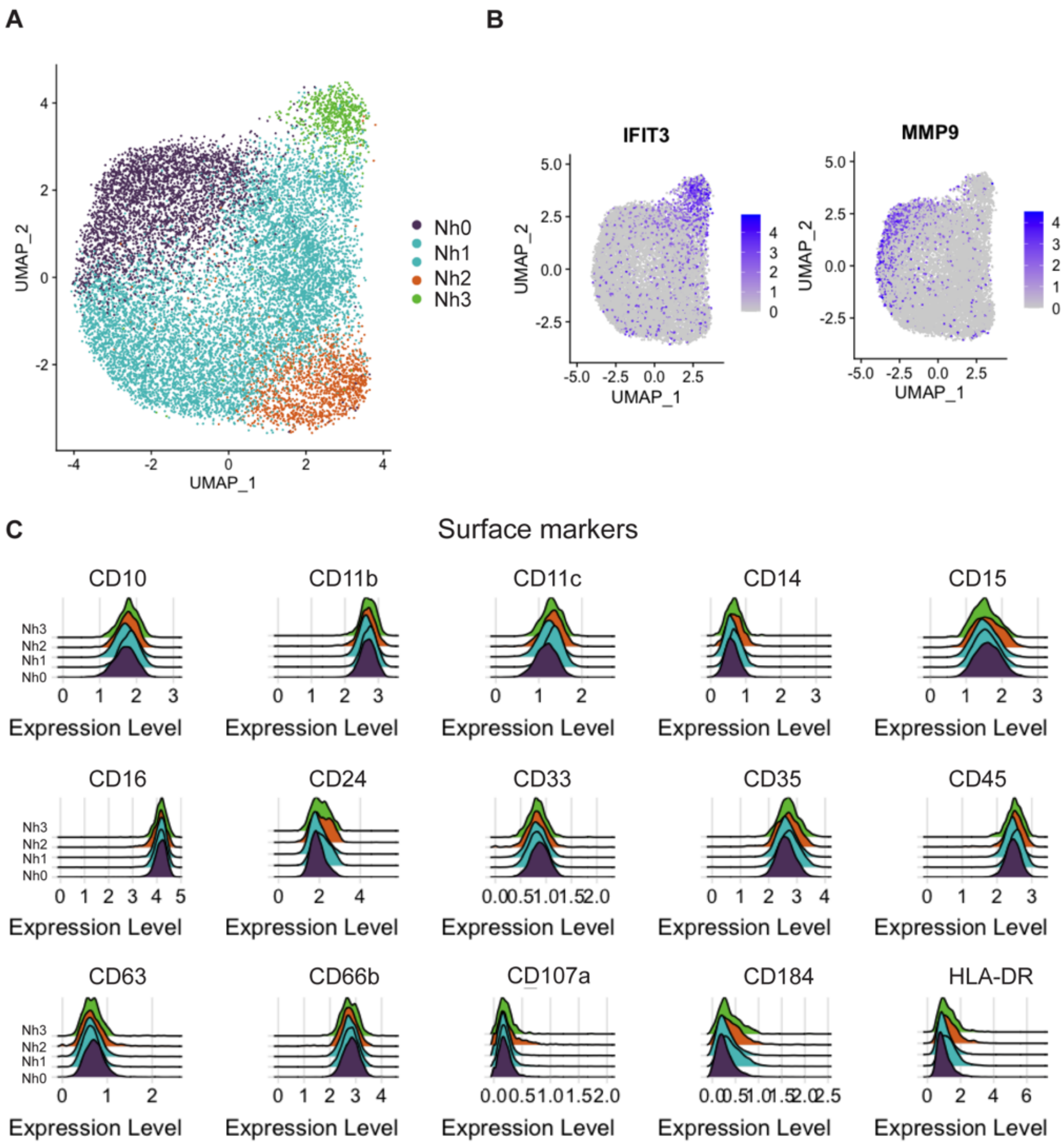


**Supplemental Figure 3**. **Surface-protein expression of neutrophil markers for the transcriptional clusters (CITE-seq). (**A) UMAP showing neutrophil transcriptional subsets from the CITE-seq analysis, colored based on clusters Nh0-Nh3 from the scRNA-seq dataset. (B) Feature plots showing representative genes for the IFN cluster (Nh3, left) and the immature cluster (Nh0, right). (C) Ridge plots showing expression of various neutrophil surface markers by transcriptional cluster.
